# KATANIN-mediated microtubule severing is required for MTOC formation and function in *Marchantia polymorpha*

**DOI:** 10.1101/2024.01.04.574198

**Authors:** Sarah Attrill, Liam Dolan

**Affiliations:** Gregor Mendel Institute, Dr Bohr-Gasse 3, 1030, Vienna, Austria; Department of Biology, University of Oxford, OX1 3RB, UK

## Abstract

Microtubule organising centres (MTOCs) are sites of localised microtubule nucleation in eukaryotic cells. Regulation of microtubule dynamics often involves KATANIN (KTN); a microtubule severing enzyme which cuts microtubules to generate new negative ends leading to catastrophic depolymerisation. In *Arabidopsis thaliana*, KTN is required for the organisation of microtubules in the cell cortex, preprophase band, mitotic spindle and phragmoplast. However, as angiosperms lack MTOCs, the role of KTN in MTOC formation has yet to be studied in plants. Two unique MTOCs – the polar organisers – form on opposing sides of the prophase nucleus in liverworts. Here we show that KTN-mediated microtubule depolymerisation is required for the *de novo* formation of polar organisers in the liverwort, *Marchantia polymorpha*. In Mp*ktn* mutants that lack KTN function, the formation, shape, number, and function of polar organisers is defective. This is in addition to defective microtubule organisation in the cell cortex, preprophase band, mitotic spindle, and phragmoplast. These data demonstrate that KTN-mediated microtubule dynamics are required for the formation of liverwort-specific MTOCs.

## Introduction

Microtubule organising centres (MTOCs) are sites of microtubule nucleation in eukaryotic cells. The centrosomes in animal cells contain centrioles which nucleate microtubules to form the mitotic spindle and astral arrays (Bettencourt-Dias and Glover, 2007). Centrioles also form in the motile spermatozoids of sperm-producing plants (bryophytes, lycophytes and monilophytes) but are not present in the somatic cells of any land plant. However acentrosomal MTOCs form in the somatic cells of bryophytes (Brown and Lemmon, 2011; Buschmann and Zachgo, 2016). In the liverwort, *Marchantia polymorpha*, two MTOCs – the polar organisers – form at opposite sides of the prophase nucleus (Brown and Lemmon, 1990). Polar organisers nucleate astral arrays that polymerise towards the cell cortex, and perinuclear arrays which encase the nucleus. In contrast to centrioles, polar organisers are not inherited but form *de novo* in each cell by aggregation of smaller microtubule nucleation sites (foci) during prophase (Buschmann *et al*., 2016). At the onset of mitosis, polar organisers disassemble and their γ-tubulin relocates to the poles of the mitotic spindle (Brown and Lemmon, 2011; Brown *et al*., 2004). Nothing is known about the proteins required for the formation or function of polar organisers.

KATANIN (KTN) catalyzes the severing of microtubules producing free negative ends which undergo catastrophic depolymerisation (McNally and Vale, 1993). By destabilising microtubules, KTN regulates microtubule organisation and dynamics (McNally and Roll-Mecak, 2018). KTN preferentially cuts microtubules at points of branching and cross-over, destabilising these discordant microtubules and generating parallel arrays in the cortex of *Arabidopsis thaliana* cells (Deinum *et al*., 2017; Zhang *et al*., 2013). In loss-of-function At*ktn* mutants, microtubules are less dynamic than in wild type cells, because the loss of microtubule severing reduces the number of free minus ends resulting in decreased rates of microtubule depolymerisation (Komis *et al*., 2017). While the defects in At*ktn* mutants demonstrate that KTN-mediated severing is required to generate dynamic microtubule arrays, it reveals nothing about the function of KTN in the formation of MTOCs. This is because, unlike bryophytes, stable MTOCs do not form in angiosperms (Chan *et al*., 2003).

We set out to test the hypothesis that KTN-mediated microtubule severing is required for the formation of MTOCs – polar organisers – in *M. polymorpha*. First, we show that the role of KTN in the organisation of interphase, mitosis and cytokinesis microtubule arrays is conserved between *M. polymorpha* and *A. thaliana*. We then demonstrate that KTN is required for the formation of polar organisers.

## Materials and Methods

### Sequence alignments and generation of phylogenetic trees

The protein sequence for the *A. thaliana KTN* p60 subunit, AT1G80350, was used in a BLASTp search against the *M. polymorpha* proteome. The top hit, Mapoly0116s0028 (Mp4g20260), was used as the query sequence for BLASTp searches in the proteome databases of 20 land plant and algal species (Table 1). In each species, except two *Osterococcus* species, sequences with E-values above E-87 were identified and selected. The protein sequences were aligned using MAFFT version 7 employing the L-INS-i method (Katoh *et al*., 2002). Sequences from three species - *Anthoceros punctatus, Salvinia cucullata* and *Picea abies* - were subsequently removed due to suspected mis-annotation of one or more exon-intron boundaries, or incomplete sequencing. For one species – *Azolla filiculoides* – a new coding sequence was proposed after suspected mis-annotation of the exon-intron boundaries, confirmed by the transcriptome. Sequences from 15 species were realigned in MAFFT and trimmed using BioEdit software to regions encoding the conserved AAA ATPase and Vsp4C domains. These domains were identified using the SMART protein domain dataset (Letunic and Bork, 2018). Maximum likelihood trees were generated using MEGA-X 10 software utilising all amino acid sites and bootstrap values were calculated from 500 replicates (Kumar *et al*., 2018).

**Table 1:**
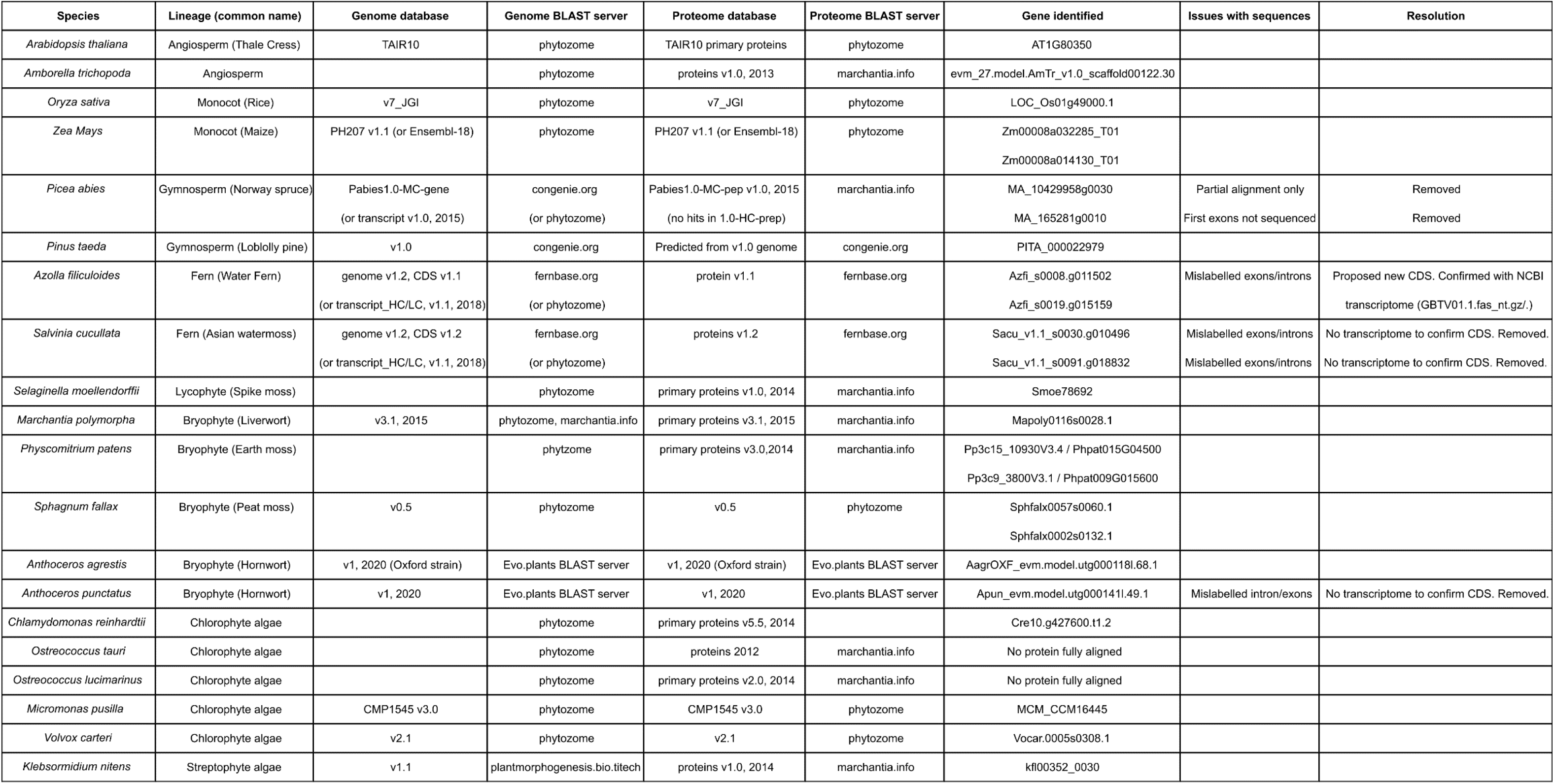
The genome and proteome databases used to identify KTN sequences in a range of land plant and algal species.

### Plant lines, growth conditions and crossings

The wild type *Marchantia polymorpha* accessions used were Takaragaike-1 (Tak-1) and Takaragaike-2 (Tak-2). All constructs were transformed into the progeny derived from crossing Tak-1 and Tak-2. Plants were grown on ½-strength B5 Gamborg’s medium containing 1.5 g/L B5 Gamborg, 0.5 g/L MES hydrate, 1% sucrose, pH adjusted to 5.5, set with 1% agar. Plants were grown at 23 °C in continuous white light at 50 - 60 μmol m²s¹.

To induce reproductive development, mature plants were potted on soil, containing a 1:3 ratio of fine vermiculite and Neuhaus N3 compost, at 20°C in long day conditions of 16 hours light, 8 hours dark. White light was set at 50 - 60 μmol m²s¹ and enhanced with far-red light at 30 - 40 μmol m²s¹. Male and female plants were crossed to generate sporangia. Sporangia were sterilised in 1 % sodium dichloroisocyanurate (NaDCC) for three minutes before washing with water and releasing the spores.

### sgRNA design for CRISPR/Cas9 mutagenesis

sgRNAs, consisting of 20 nucleotides followed by a NGG sequence, were designed to target the Mp*KTN* (Mapoly0116s0028/Mp4g20260) gene using the CRISPR-P software (Lei *et al*., 2014). Candidate sequences were checked for off-target hits by a BLAST search against the *M. polymorpha* genome v4 (marchantia.info). The two final sequences were sgRNA-K3 TACGTTGGCCTCCAAATGGA(GGG) – to which a CTCG overhang was added to the 5’ end prior to cloning - and sgRNA-K5 GGAGCTTGCCAGACGTACAG(AGG).

### Cloning of CRISPR/Cas9 plasmids

Cloning of the sgRNA-K3 Cas9 plasmid used the vectors and protocol presented in Thamm *et al*., 2020 adapted from Sugano *et al*., 2014. Cloning of the sgRNA-K5 Cas9 plasmid was performed by the Protein Technologies Facility at Vienna BioCenter Core Facilities using vectors from the OpenPlant toolkit and following the protocol published in Sauret-Güeto *et al*., 2020.

### Transformation of constructs into *M. polymorpha*

Plasmids were transformed into *M. polymorpha* sporelings using transgenic *Agrobacterium* following the method developed in Ishizaki *et al*., 2008 and improved upon in Honkanen *et al*., 2016. Transgenic plants were selected by their resistance to 10 µg/mL hygromycin.

### DNA extraction and sequencing of *M. polymorpha*

DNA extraction and amplification used the Phire Plant Direct PCR kit (ThermoFisher Scientific) (PCR primers, Table 2) and GeneJet Gel Purification kit (ThermoFisher Scientific). Alternatively, DNA was extracted from 3 x 3 mm pieces of plant tissue ground within 100 μL of extraction buffer; 100 mM Tris HCl, pH 9.5, 1 M KCl, 10 mM EDTA. Samples were incubated at 65 °C for 10 minutes before dilution in 500 µL MonoQ water. DNA was PCR amplified using 2x HS Taq Polymerase; 2x Hot Start MM, 0.5 µM forward PCR primer, 0.5 µM reverse PCR primer, 1 µL DNA, 7 µL Nuclease free water. The PCR products were purified using an the ExoSAP mix; 0.04 µL Exonuclease I, 0.4 µL Shrimp Alkaline Phosphatase, 1.56 µL Storage solution. The reaction was incubated at 37 °C for 30 minutes, before deactivation at 80 °C for 10 minutes.

**Table 2:**
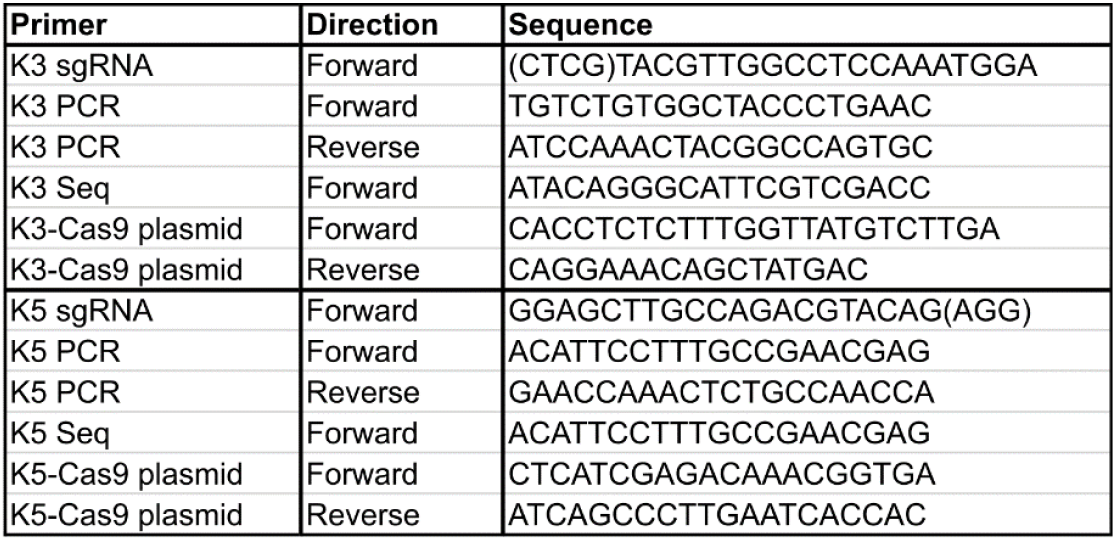
sgRNA sequences used to target MpKTN for mutagenesis, primers for sequencing MpKTN gene and primers to test for Cas9 presence in transformed plants.

Isolated DNA was Sanger sequenced (Seq primers in Table 2). Sequences were aligned against the Tak-1 genome sequence using Geneious or CLC Genomics Workbench to identify mutations at the PAM site.

### Generation and selection of CRISPR/Cas9 mutants expressing reporters

Mp*katanin* (Mp*ktn*) mutants were crossed to wild type plants expressing the microtubule reporter, *p*Mp*EF1α:GFP-*Mp*TUB1* (GFP-MpTUB1), generated by Buschmann *et al*., 2016. The resulting F1 progeny were grown, genotyped for mutations (PCR primers in Table 2), and tested for the presence of the Cas9 plasmid (Cas9 primers in Table 2). Cas9-free lines were screened for GFP-MpTUB1 by testing for resistance to 10 µg/mL hygromycin and fluorescence imaging. Cas9-free wild type (wildtype-TUB) and Mp*ktn* (Mp*ktn*-TUB) siblings expressing the GFP-MpTUB1 reporter were selected (SFig. 1F).

**Figure 1:**
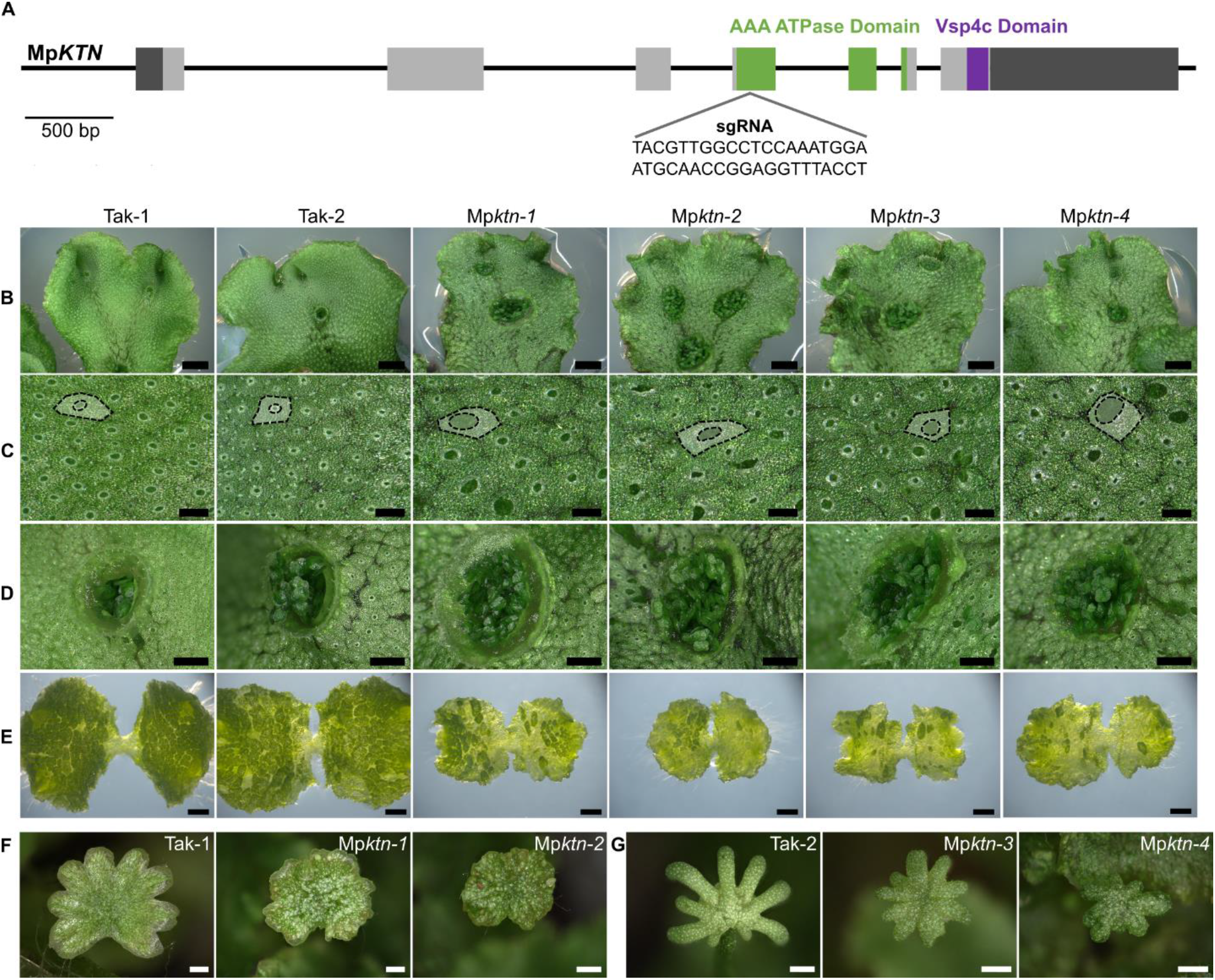
KTN is required for tissue and organ development in *M. polymorpha*. **(A)** Schematic of the Mp*KTN* gene indicating one sgRNA location and sequence. Light grey boxes indicate exons and dark grey boxes indicate untranslated regions. Green indicates regions encoding the AAA ATPase domain. Purple indicates the region encoding the Vsp4c domain. Scale bar, 500 bp. **(B)** Dorsal thallus of Tak-1, Tak-2 and four Mp*ktn* mutants. Scale bars, 2 mm. **(C)** Dorsal epidermal surface of Tak-1, Tak-2 and four Mp*ktn* mutants. An air chamber and air pore are outlined and false coloured in each image. Scale bars, 300 μm. **(D)** Gemma cups on the dorsal thallus of Tak-1, Tak-2 and four Mp*ktn* mutants. Scale bars, 1 mm. **(E)** 9-day-old gemmalings from Tak-1, Tak-2 and four Mp*ktn* mutants. Scale bars, 1 mm. **(F)** Antheridiophores of Tak-1 and two male Mp*ktn* mutants grown under far-red light. Scale bars, 1 mm **(G)** Archegoniophores of Tak-2 and two female Mp*ktn* mutants grown under far-red light. Scale bars, 1 mm.

### Stereomicroscope imaging of plant tissues

Gemmalings were imaged with the Leica MZ16FA stereomicroscope equipped with the Leica DFC300 FX camera. Mature plants and tissues were imaged with the Keyence VHX-7000 digital equipped with a VHX-7020 camera and VH-Z00R/T and VH-ZST lenses.

### Spinning disk imaging of microtubules

Imaging chambers were adapted from Kirchhelle and Moore, 2017. A breathable gum boarder (Carolina Observation gel) was filled with Gamborg media and layered with cellophane soaked in liquid Gamborg media (½-strength B5 Gamborg’s medium without agar). Perfluorodecalin and 0-day gemmae were added, and the chamber sealed with a cover slip. Gemmae were grown for 2 days within the chamber prior to imaging.

Microtubules were imaged with an Olympus IX3 Series (IX83) inverted microscope equipped with a Yokogawa W1 spinning disk, Hamamatsu ORCA-Fusion CMOS camera and a 100x/1.45 NA oil objective. Samples were excited at 488 nm and emission captured at 525 nm. Z-stacks were taken with 0.26 µm slices. GFP-MpTUB1 labelled cortical arrays were imaged in the central epidermal cells, and mitotic arrays in dividing cells near the meristem.

### Deconvolution and conversion of spinning disk images

Image deconvolution used Huygens software (Scientific Volume Imaging). Z-projections and central slices were converted using ImageJ Fiji (Schindelin *et al*., 2012). For temporal projections, each image slice was first deconvolved in Huygens then maximum projections were generated for each timepoint in ImageJ Fiji before temporal projection.

### Analysis of microtubule organisation

Cortical microtubule were analysed using the ImageJ LPX package published in Higaki *et al*., 2010 following the steps in Higaki, 2017. From Z-projections of the central epidermis, individual cells were outlined using the Freehand Tool in ImageJ Fiji. Microtubules were skeletonised using the LPX Filter2d with the Otsu method and a line extract value of 5. The image was masked by the cell outlines to identify the skeletonised microtubules in each cell for analysis using the LPX script. Statistical analysis used Microsoft Excel and graphs were made in R.

The number of polar organisers per cell were manually counted from Z-projections. Spindle length was measured in ImageJ Fiji using the Line tool from Z-projections. Spindle position was quantified by identifying the central point of the cell and spindle in ImageJ Fiji then calculating the distance between these points. Statistical analysis used Microsoft Excel.

## Results and Discussion

### There is a single *KTN* gene in *Marchantia polymorpha* and mutations in Mp*KTN* result in defective organ development

To define the function of KTN in polar organiser formation, we characterised the phenotype of mutants carrying loss of function mutations in the Mp*KTN* gene. To identify *KTN* genes in *Marchantia polymorpha,* the protein sequence for the *A. thaliana KTN* p60 subunit, AT1G80350, was used as a query in a BLASTp search against the *M. polymorpha* proteome. Mapoly0116s0028 (Mp4g20260) was the most similar sequence identified. Two phylogenetic trees were then constructed using KTN protein sequences from a variety of land plant and algal species (SFig. 1A, B). These sequences were identified using the *M. polymorpha* sequence as a query in BLASTp searches across multiple databases (Table 1). The topology of the two trees indicates that Mapoly0116s0028 (Mp4g20260) is a KTN family member and is the only *KTN* p60 subunit gene in *M. polymorpha* (SFig. 1A, B). This gene will now be referred to as Mp*KTN*.

To investigate the function of KTN in polar organiser formation, we used CRISPR/Cas9 mutagenesis to generate loss of function mutations in Mp*KTN*. sgRNAs were designed to target regions in the Mp*KTN* gene encoding for the AAA ATPase catalytic domain and Vsp4C domain (Fig. 1A; SFig. 1C). Mutations in these regions of the *KTN* gene in *A. thaliana* result in a loss of KTN function (Luptovčiak *et al*., 2017). A series of Mp*KTN* mutant alleles – including nucleotide deletions, insertions, and substitutions – were produced that altered the predicted amino acid sequence by inducing frameshifts, substitutions, and protein truncations (SFig. 1D, E). All Mp*ktn* mutants developed similar phenotypes with multiple developmental defects compared to wild type. This included a crinkled thallus, enlarged air chamber pores, enlarged gemma cups, irregular gemmaling development, delayed plant growth, and abnormal reproductive organs (Fig. 1B-G; SFig. 2, 3). Many of the developmental defects reflect those in At*ktn* (Luptovčiak *et al*., 2017). The consistent defective phenotype of all Mp*ktn* mutants, in combination with the mutations being located within regions encoding highly conserved functional domains, suggests that Mp*ktn* have a complete loss of KTN function.

**Figure 2:**
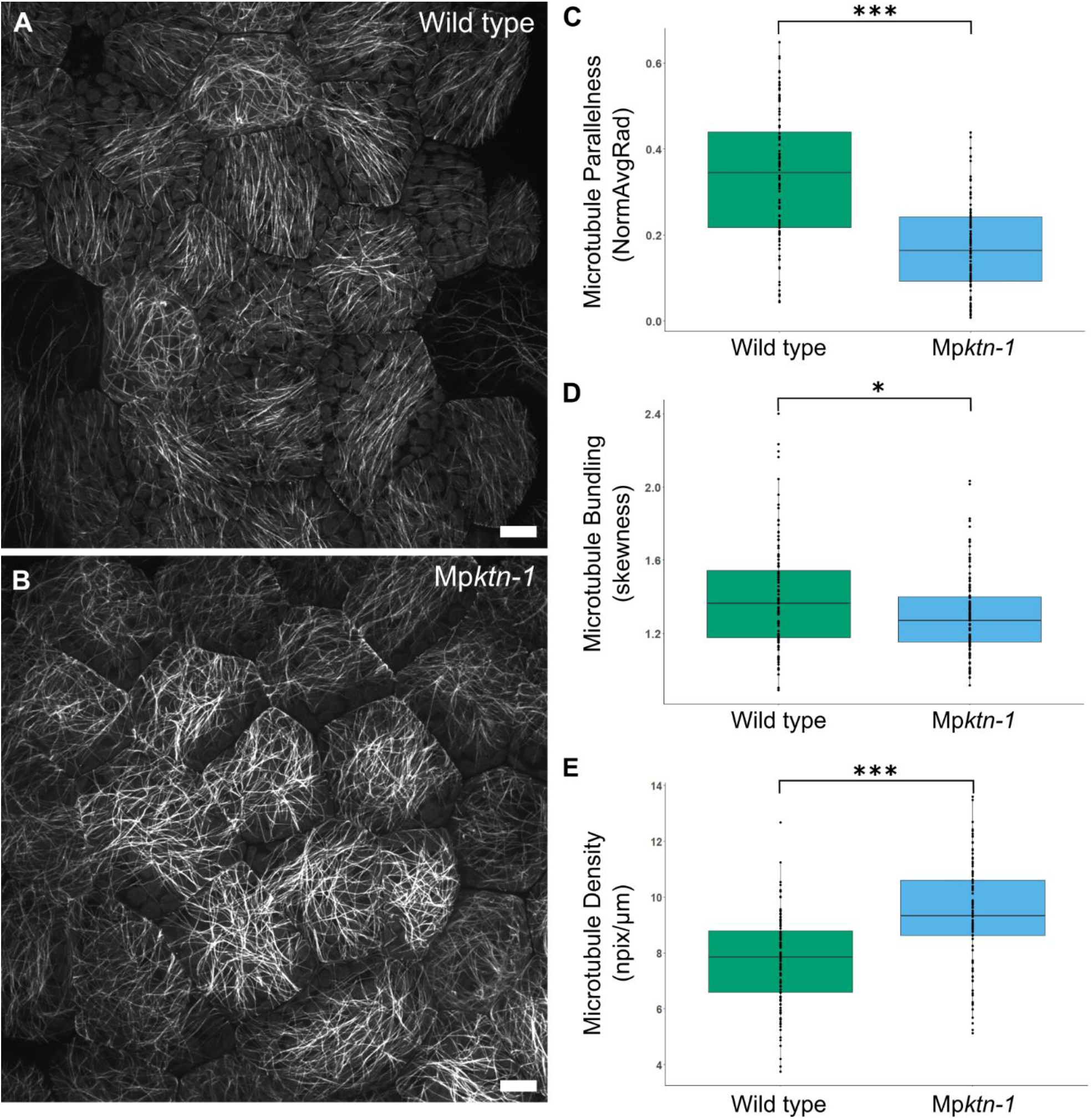
KTN promotes parallelness and bundling of cortical microtubules. **(A, B)** Cortical microtubule arrays in central epidermal cells of wild type (A) and Mp*ktn-1* (B) gemmalings. Presented are Z-projections. Scale bars, 10 μm. **(C - E)** Boxplots of the parallelness (C), bundling (D), and density (E) of cortical microtubules in wild type and Mp*ktn-1* epidermal cells. N = 108 cells for wild type, N = 107 cells for Mp*ktn-1* across 10 gemmalings per genotype. Data was analysed using Welches 2-tail T-test and 2-tail F-test, see SFig. 4A. Significant differences in the T-test value between the genotypes is indicated by (*) P ≤ 0.05, (**) P ≤ 0.01 and (***) P ≤ 0.001.

### KTN activity is required for the organisation of cell-specific cortical arrays in the gemma epidermis

KTN-mediated microtubule severing is required for the organisation of microtubules in the cortex of *A. thaliana* cells (Komis *et al*., 2017). To test if KTN controls microtubule organisation in *M. polymorpha,* the arrangement of cortical arrays in Mp*ktn* was investigated. Mp*ktn-1* and Mp*ktn-2* – harbouring mutations in the Mp*KTN* gene at the AAA ATPase-encoding domain – were crossed to wild type (Mp*KTN*) plants expressing the microtubule reporter, *p*Mp*EF1α:GFP-*Mp*TUB1* (Buschmann *et al*., 2016). Cas9-free wild type and Mp*ktn* expressing GFP-MpTUB1 were selected from the F1 progeny (SFig. 1F). Microtubules in the cortex of the outer periclinal wall of the epidermal cells at the centre of 2-day-old gemmalings were imaged. In wild type, two arrangements of cortical arrays were observed. Some cells developed parallel arrays, while other cells developed non-parallel arrays (Fig. 2A; SFig. 4B). By contrast, cortical arrays were consistently randomly organised in all cells of Mp*ktn-1* and Mp*ktn-2* (Fig. 2B; SFig. 4C). Quantification revealed significant differences between wild type and Mp*ktn-1*. Microtubule parallelness and bundling were on average lower in Mp*ktn-1* than in wild type, but with higher variation between cells (Fig. 2C, D; SFig. 4A). While microtubule density was consistently greater in Mp*ktn-1* cells than in wild type (Fig. 2E; SFig. 4A). These data indicate that KTN-mediated microtubule depolymerisation is required for the organisation of cortical microtubules into parallel, bundled, low-density arrays in *M. polymorpha*. Further, these data demonstrate that MpKTN functions in a cell-specific manner to generate variation in the microtubule organisation between individual epidermal cells.

### KTN is required for the formation of two, compact polar organisers per cell

Since microtubule organising centres (MTOCs) – sites of localised microtubule nucleation – are not present in *A. thaliana*, the role of KTN-mediated microtubule severing in MTOC formation has not been established in land plants (Luptovčiak *et al*., 2017). However, two MTOCs, known as polar organisers, form *de novo* at opposite poles of the prophase nucleus in *M. polymorpha* (Brown and Lemmon, 2011; Buschmann *et al*., 2016). Polar organisers nucleate astral microtubules that polymerise into the cell cortex, and perinuclear microtubules that polymerise towards the nucleus equator (Fig. 3A). We hypothesised that KTN-mediated microtubule severing would be required for the formation and organisation of polar organisers.

**Figure 3:**
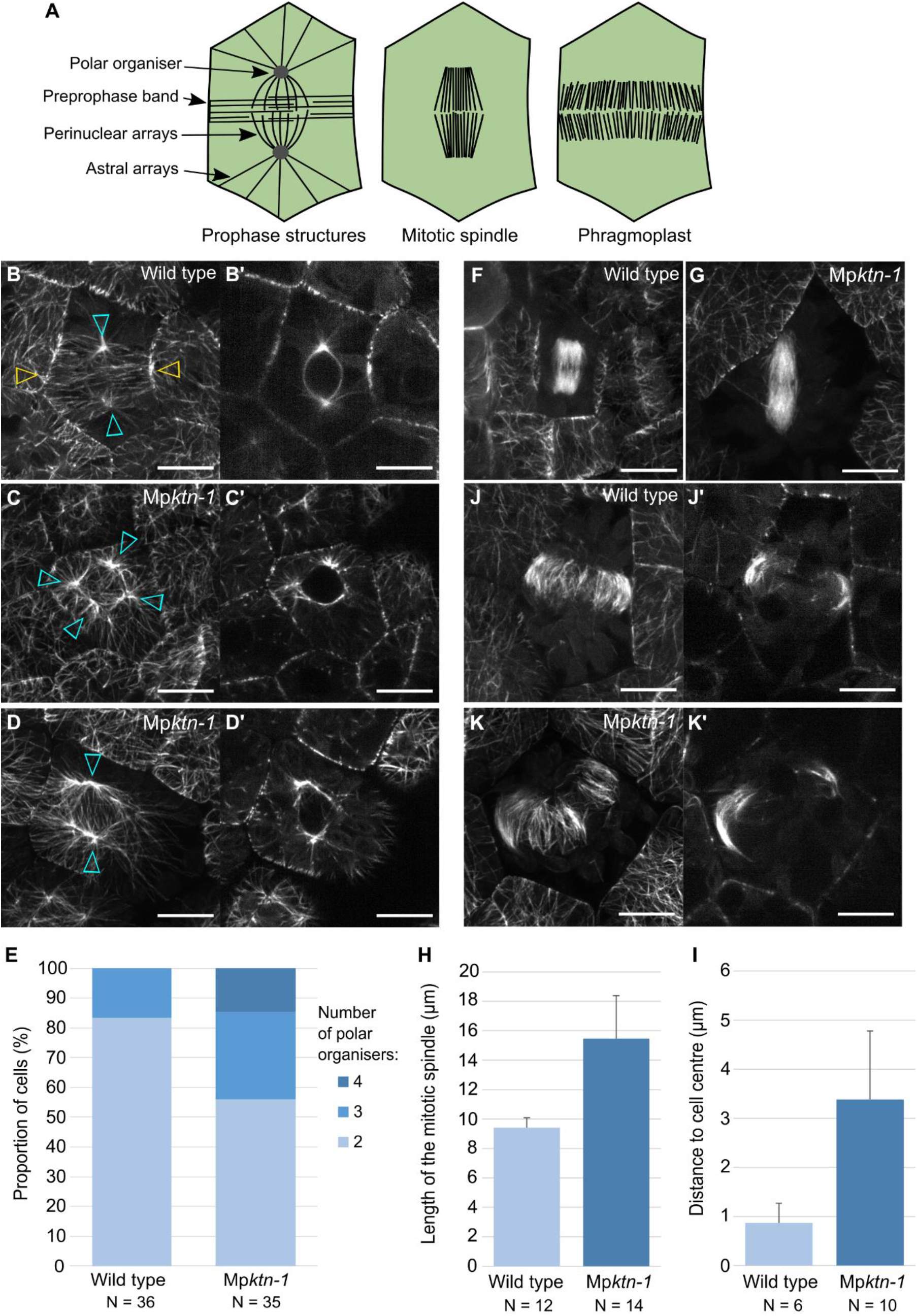
KTN regulates the number and organisation of polar organisers in each cell. **(A)** Microtubule organisations in *M. polymorpha* cells during prophase, mitosis, and cytokinesis. **(B - D)** Polar organisers and astral arrays in wild type (B) and two Mp*ktn-1* (C, D) cells. Presented are Z projections (B-D) and central slices in XY (B’-D’). Scale bars, 10 μm. Cyan arrows indicate polar organisers, yellow arrows indicate the preprophase band. **(E)** Proportion of wild type and Mp*ktn-1* cells with 2, 3 or 4 polar organisers. N = 36 cells for wild type, and N= 35 cells for Mp*ktn-1*. **(F, G)** Mitotic spindle in wild type (F) and Mp*ktn-1* (G) cells. Presented are Z-projections. Scale bars, 10 μm. **(H)** Graph of the mitotic spindle length (μm) in wild type and Mp*ktn-1*. N is the number of spindles. **(I)** Graph of the distance (μm) between the centre of the mitotic spindle and the centre of the cell for wild type and Mp*ktn-1.* N is the number of spindles. **(J, K)** The phragmoplast in wild type (J) and Mp*ktn-1* (K) cells. Presented are Z-projections (J, K) and central slices in XY (J’, K’). Scale bars, 10 μm.

To test the hypothesis that KTN is required for polar organiser formation, the microtubules in 2-day-old gemmalings from wild type and Mp*ktn-1* were imaged. Wild type formed two distinct polar organisers at opposite sides of the nucleus (Fig. 3B). By contrast, Mp*ktn-1* often formed more than two polar organisers resulting in a multipolar structure around the nucleus (Fig. 3C, D). Quantification showed that wild type had two, or occasionally three, polar organisers per cell (Fig. 3E). By contrast, Mp*ktn-1* had up to four distinct polar organisers per cell. In wild type, the astral microtubules radiated symmetrically towards the cell cortex and the perinuclear microtubules formed a bipolar array (Fig. 3B). By contrast, both astral and perinuclear arrays were denser and relatively disorganised in Mp*ktn-1* (Fig. 3C, D). Furthermore, the astral microtubules radiated asymmetrically to the cell cortex. Overall these data indicate that KTN-modulated microtubule severing is required for the formation of two polar organisers at opposing sides of the nucleus.

The preprophase band (PPB) is a parallel microtubule array that forms in the cell cortex during prophase – after polar organiser formation – and predicts the future cell division plane (Brown and Lemmon, 1990; Rasmussen *et al*., 2013). A PPB formed in most wild type cells with two polar organisers (Fig. 3B). In Mp*ktn-1* cells with polar organisers, no distinctive PPB formed but randomly organised microtubules were present in the cell cortex (Fig. 3C, D). This suggests that the initial steps of PPB formation are KTN-dependent, and that the formation of a dense, parallel microtubule array requires MpKTN.

To investigate the role of KTN in mitotic spindle and phragmoplast formation, microtubules in dividing wild type and Mp*ktn-1* cells were imaged. Mitotic spindles were short, box-shaped, and centrally positioned in wild type cells (Fig. 3F). By contrast, the mitotic spindles were significantly longer and with tapered ends in Mp*ktn-1* (Fig. 3G, H). Similar elongated spindles were observed when KTN was inhibited or knockdown in *Caenorhabditis elegans* embryos, mouse oocytes and *Xenopus tropicalis* cells (Gao *et al*., 2019; Loughlin *et al*., 2011; McNally *et al*., 2006). Mitotic spindles were also positioned significantly further from the cell center in Mp*ktn-1*, but with greater variability, than in wild type (Fig. 3I). Phragmoplasts in wild type extended across the cell to divide the cell into two relatively equal parts (Fig. 3J). By contrast, phragmoplasts in Mp*ktn-1* were often bent along their length and positioned to divide the cell into two unequal parts (Fig. 3K). We conclude that KTN regulates mitotic spindle length and position, as well as phragmoplast morphology and position in *M. polymorpha*.

Overall, we conclude that KTN is required for the formation and integrity of the polar organisers and the PPB in *M. polymorpha*. Polar organisers form by aggregation of smaller foci around the nucleus into two bipolar centres, as described in Buschmann *et al*., 2016. Our data are consistent with the hypothesis that KTN-mediated severing, and the resulting catastrophic depolymerisation of microtubules, is required for the aggregation of multiple foci into two polar organisers. We further hypothesise that KTN-mediated severing is similarly required for the reorganisation of cortical microtubules into a PPB.

### KTN controls the position of the new cell plate by aligning the mitotic spindle axis with the polar organiser axis

It has been proposed that the PPB and polar organisers together determine the position of the mitotic spindle, phragmoplast and ultimately the plane of cell division in *M. polymorpha* (Buschmann *et al*., 2016). We hypothesised that in Mp*ktn*, the disordered polar organisers were the origin of the mispositioned mitotic spindles and phragmoplasts. This would result in abnormal cell division planes in Mp*ktn*. To investigate if polar organisers orient the cell division plane, timelapse imaging of microtubule organisation in dividing wild type and Mp*ktn-1* cells was performed.

We compared the relationship between the orientation of the polar organiser axis (the axis between the two polar organisers), the mitotic spindle axis (the axis between the two spindle poles), and the plane of phragmoplast expansion in each cell. In wild type, the polar organisers and their associated bipolar perinuclear arrays were positioned near the cell centre (0 to 30 minutes in Fig. 4A). The polar organiser axis was parallel to the cell surface and perpendicular to the PPB. Later, the mitotic spindle formed in the cell centre with an axis parallel to the polar organiser axis and remained in this orientation throughout mitosis (40 to 50 minutes in Fig. 4A). These observations of polar organiser dynamics during wild type cell division were identical to those previously described in Buschmann *et al*., 2016. In Mp*ktn-1*, it was difficult to define the polar organiser axis as there was often supernumerary polar organisers and a disorganised, multipolar perinuclear array. We therefore defined the polar organiser axis as a line between the two brightest microtubule foci. In Mp*ktn-1*, the polar organiser axis was often tilted within the cell e.g. not parallel to cell surface as defined by the anticlinal cell walls (cell walls oriented perpendicular to the epidermis surface) (0 to 40 minutes in Fig. 4B, C). When the mitotic spindle formed, its axis was oblique relative to the polar organiser axis (clearly viewed in the XZ plane at 50 to 80 minutes in SFig. 5D). Furthermore, the mitotic spindle elongated and rotated overtime in Mp*ktn-1* (60 to 110 minutes in Fig. 4C). This indicates that the spatial relationship between the polar organiser axis and the mitotic spindle axis are uncoupled in Mp*ktn-1*.

**Figure 4:**
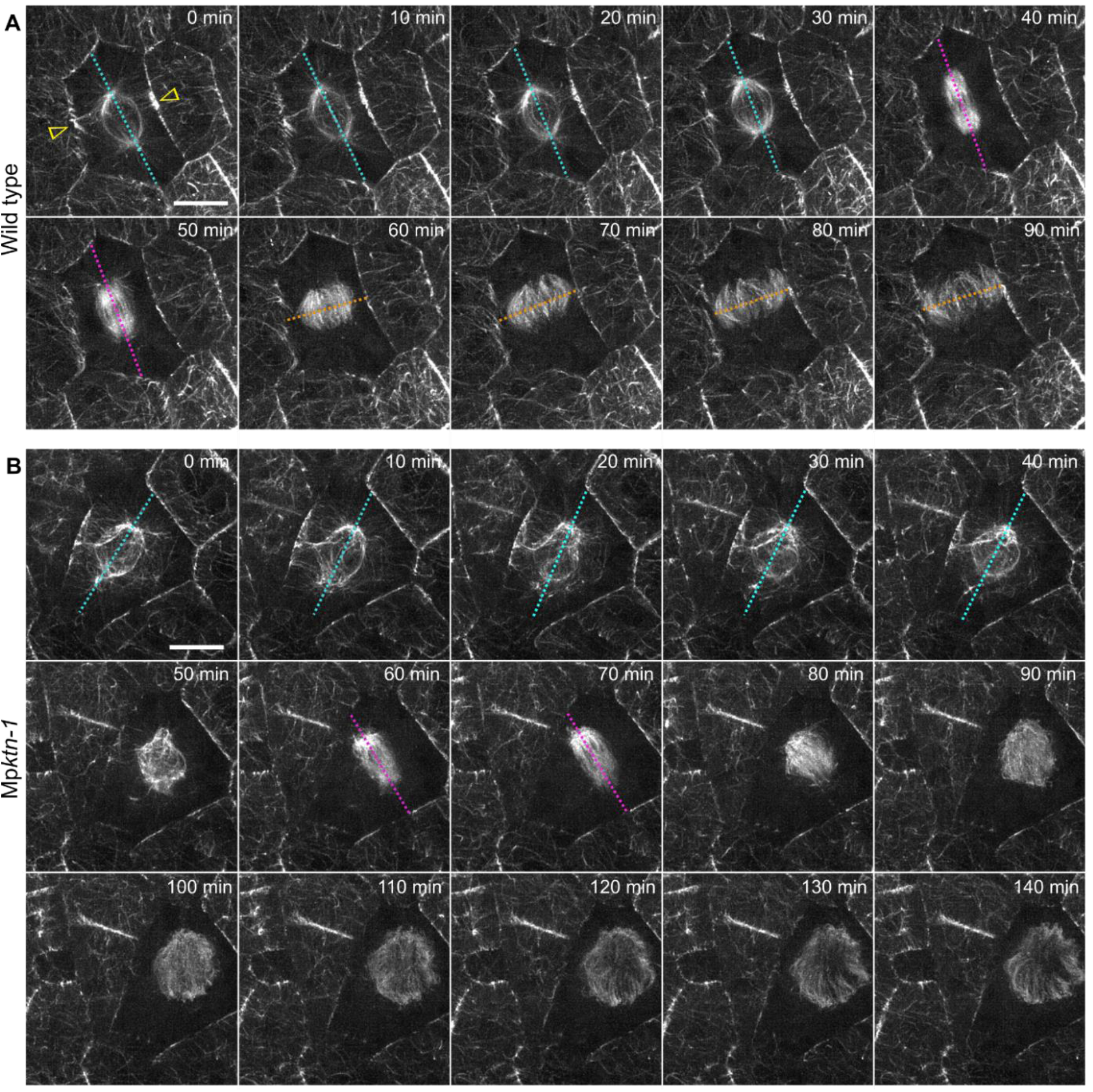
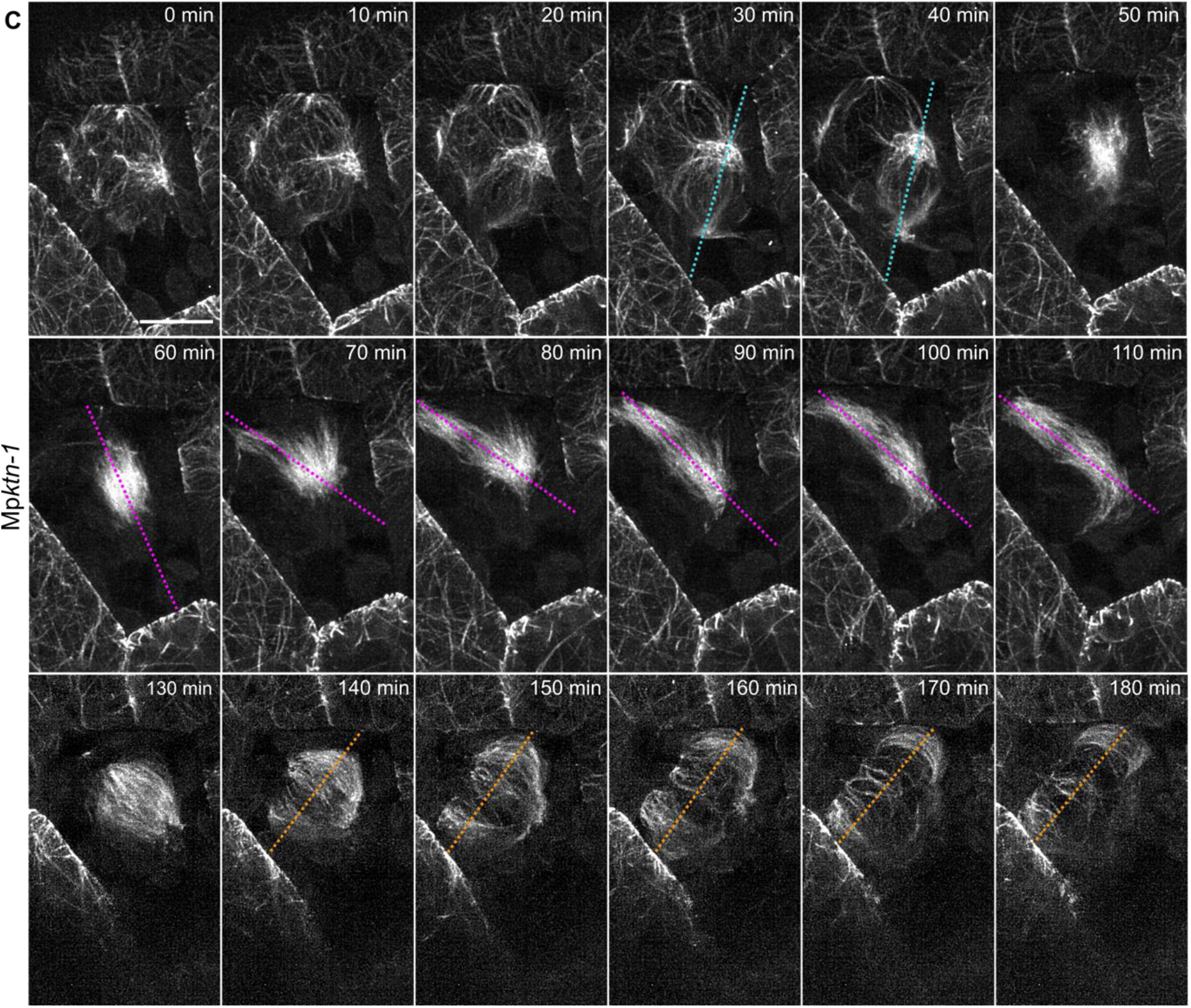
The polar organiser axis and mitotic spindle axis misalign in dividing Mp*ktn-1* cells. Timelapses of microtubules in dividing wild type **(A)** and Mp*ktn-1* **(B, C)** cells from 2-day-old gemmae. Presented are deconvolved Z-projections taken at 10-minute intervals. Dotted cyan lines indicate the polar organiser axis, dotted magenta lines indicate the mitotic spindle axis and dotted orange lines indicate the phragmoplast axis. Yellow arrows indicate the preprophase band. Scale bars, 10 μm.

**Figure 5:**
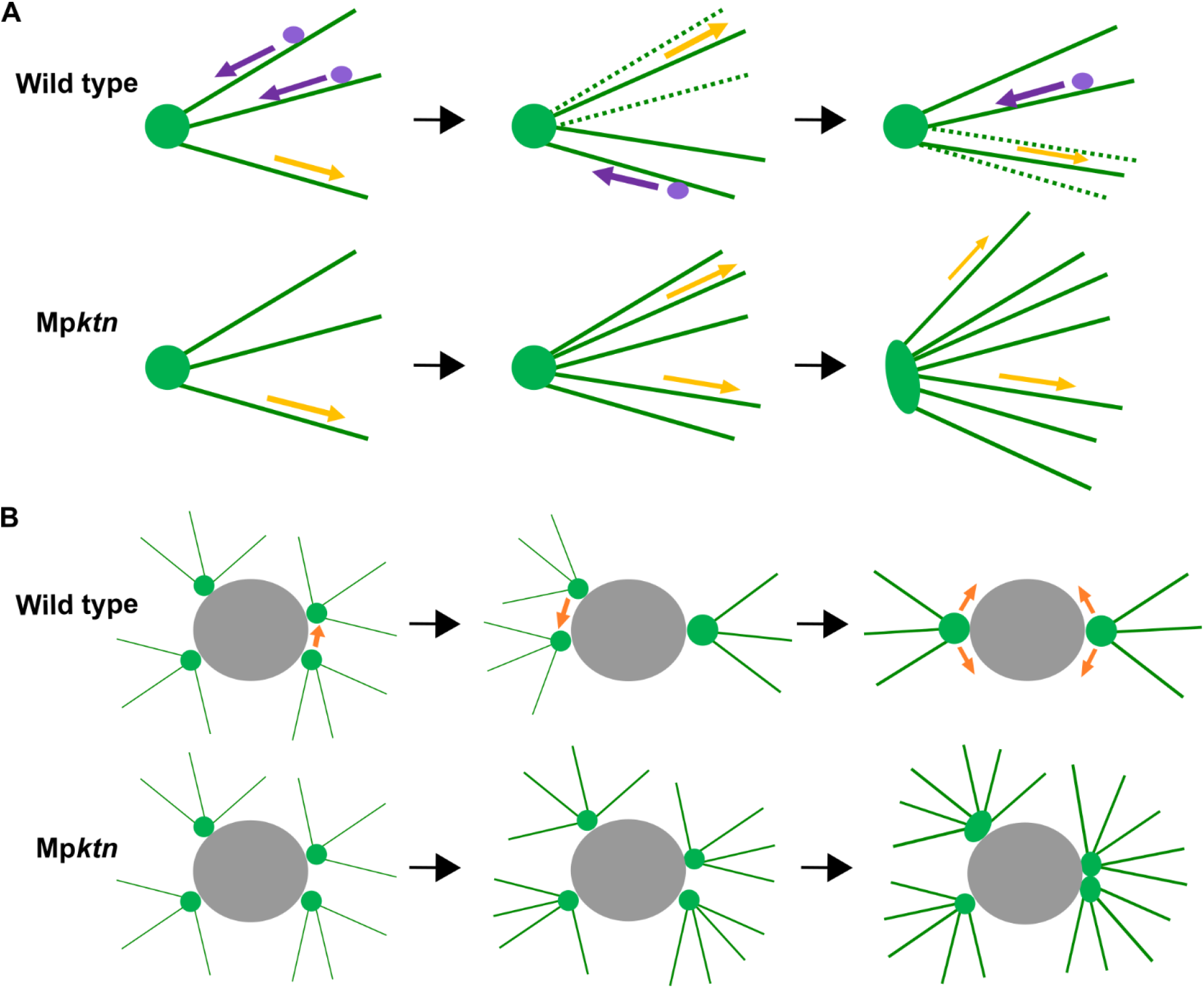
Hypotheses for the role of KTN in astral microtubule dynamics and the formation of polar organisers in *Marchantia polymorpha*. **(A)** Hypothesis for the dynamics of astral microtubules in wild type and Mp*ktn*. Astral microtubules continuously polymerise from polar organisers. In wild type, KTN cuts some astral microtubules resulting in their depolymerisation. Astral array density remains stable over time. In Mp*ktn*, no astral arrays are severed and depolymerised. Astral array density increases over time and the polar organiser expands to accommodate. Green lines represent microtubules, green circles represent polar organisers and purple dots represent KTN proteins. Yellow arrows indicate microtubule polymerisation and purple arrows indicate microtubule depolymerisation after KTN severing. **(B)** Hypothesis for the formation of polar organisers in wild type and Mp*ktn*. Multiple microtubule nucleation sites (foci) form around the nucleus in prophase. In wild type, these foci can move and fuse to form two polar organisers. In Mp*ktn*, these foci cannot move and fuse due to their stable astral arrays. Each foci becomes a polar organiser, leading to a multipolar structure. Green lines represent microtubules, green circles represent microtubule foci and polar organisers, and grey circles represent the nucleus. Orange arrows indicate the movement of microtubule foci and polar organisers.

The phragmoplast formed and expanded in a plane perpendicular to the last position of the mitotic spindle in both wild type and Mp*ktn-1* cells. In wild type, the phragmoplast formed in the cell centre and expanded in the same plane as the PPB (60 to 90 minutes in Fig. 4A). The new cell wall formed perpendicular to the surface plane of the cell – anticlinal – and split the cell into two more or less equal halves i.e., division was symmetric (90 minutes in SFig. 5A). By contrast in Mp*ktn-1*, as the mitotic spindle was not oriented parallel cell surface and/or rotated before cytokinesis, the phragmoplast divided the cell into two unequal parts (140 to 180 minutes in Fig. 4C) or oblique relative to the surface plane of the cell (clearly viewed in the ZY plane at 90 to 140 minutes in SFig. 5B). These data indicate that KTN-mediated microtubule severing is required to orient the mitotic spindle and the subsequent phragmoplast for symmetrical, anticlinal divisions.

Taken together, our data demonstrate that KTN-mediated microtubule severing is required for the *de novo* formation of MTOCs in *M. polymorpha*. This is a novel function for KTN in land plants. We also show that severing of microtubules by KTN, leading to microtubule depolymerisation, is required for organisation of cell-specific cortical arrays. We further show that KTN is required to orient cell divisions through its functions in PPB formation, MTOC positioning, and mitotic spindle alignment and stabilisation. Roles for KTN in the dynamic reorganisation of cortical and mitotic microtubule arrays have previously been described in the angiosperm, *A. thaliana* (Lindeboom *et al*., 2013; Komis *et al*., 2017). However, the role of KTN in plant MTOC formation and function has never been described.

Bryophytes develop MTOCs – localised sites of microtubule nucleation – unlike other land plants that develop delocalised sites of microtubule polymerisation (Chan *et al*., 2003; Brown & Lemmon, 2011). Polar organisers are liverwort-specific MTOCs that contain γ-tubulin and nucleate two distinct populations of microtubules: the astral arrays and the perinuclear arrays (Brown *et al*., 2004, 2011; Buschmann *et al*., 2016). In loss of function Mp*ktn* mutants, polar organisers were generally larger, more elongated, and nucleated denser arrays than in wild type cells. We therefore speculate that KTN-mediated microtubule depolymerisation is required to restrict the size of each polar organiser (Fig. 5A). This is consistent with the hypothesis that the polar organiser size is proportional to the number of attached microtubules.

We also speculate that KTN-mediated microtubule depolymerisation is required for the development of bipolar pairs of polar organisers. Two polar organisers form in each wild type cell through the fusion of microtubule foci in a 3-hour period before mitosis - a process called bipolar aggregation by Buschmann *et al.,* 2016. We show that supernumerary polar organisers form in Mp*ktn* cells. This indicates that without functional KTN, the number of microtubule foci often did not reduce to two before mitosis. If the bipolar aggregation model proposed by Buschmann *et al*. is correct, our data would be consistent with the hypothesis that KTN-mediated microtubule depolymerisation is required for the aggregation and/or fusion of microtubule foci into polar organisers (Fig. 5B).

Together, these data demonstrate that KTN-mediated microtubule dynamics is required for the *de novo* formation of MTOCs in *M. polymorpha*, a novel function for KTN in land plants.

## Supplementary Data

**Supplementary Figure 1:**
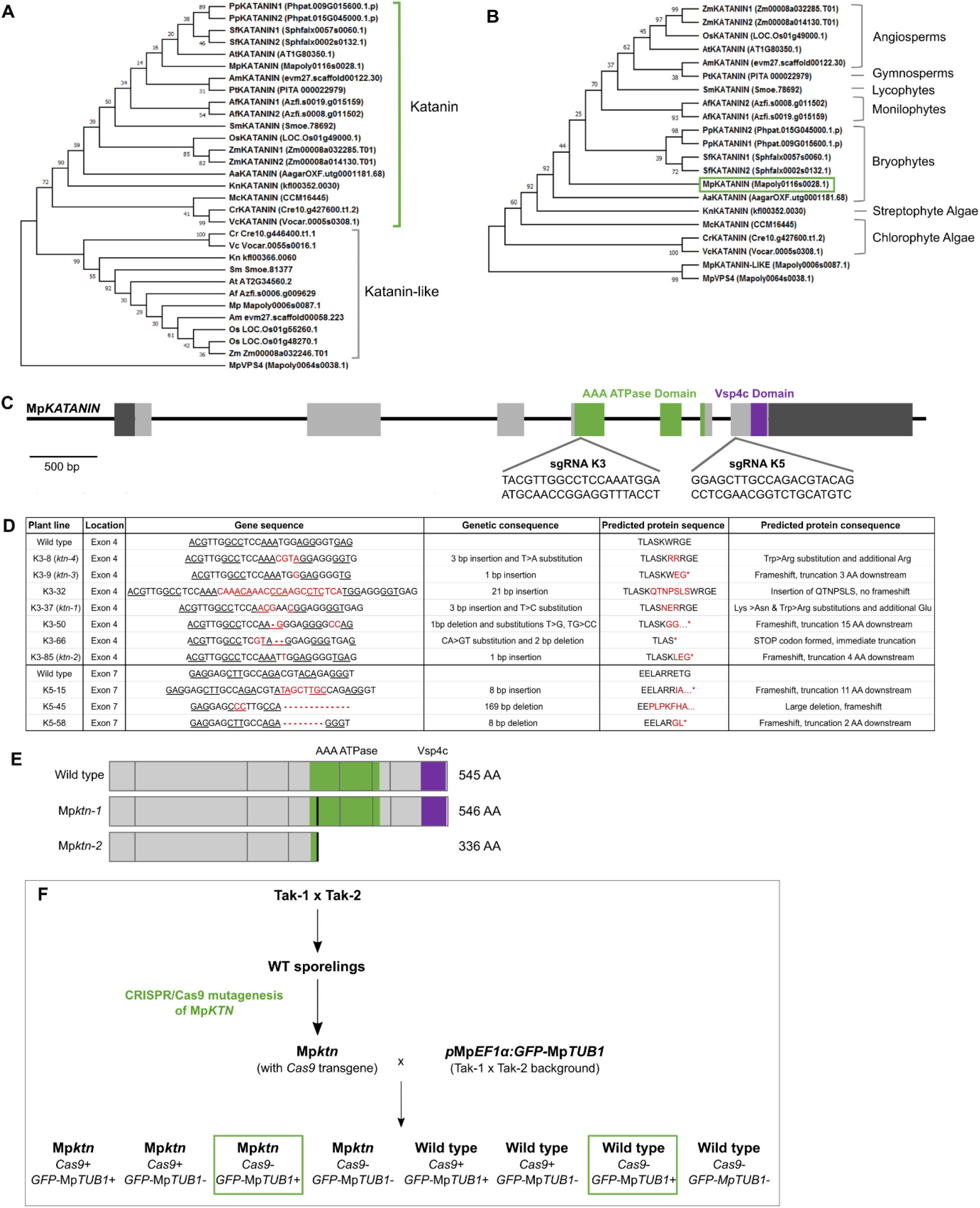
*Marchantia polymorpha* has a single *KTN* gene. **(A)** Maximum likelihood tree of the KTN and KTN-LIKE proteins from 15 land plant and algal species, based on full protein sequence alignments. Includes MpVSP4 (Mapoly0064s0038) protein as an outgroup. The KTN and KTN-like clades are labelled. **(B)** Maximum likelihood tree of KTN protein from 15 plant and algal species, based on the aligned AAA domain of protein sequences. Includes MpVSP4 (Mapoly0064s0038) and MpKTN-LIKE (Mapoly0006s0087) proteins. The major plant lineages are labelled and the single Mp*KTN* (Mapoly0116s0028) is indicated by a green box. At, *Arabidopsis thaliana.* Am, *Amborella trichopoda.* Os, *Oryza sativa.* Zm, *Zea mays*. Pt, *Pinus taeda.* Af, *Azolla filiculoides.* Sm, *Selaginella moellendorffi.* Mp, *Marchantia polymorpha.* Pp, *Physcomitrium patens.* Sf, *Sphagnum fallax.* Aa, *Anthoceros agrestis*. Cr, *Chlamydomonas reinhardtii.* Mc, *Micromonas pusilla.* Vc, *Volvox carteri*. Kn, *Klebsormidium nitens*. **(C)** Schematic of the Mp*KTN* gene indicating the location and sequences of the two sgRNAs (K3 and K5). Light grey boxes indicate exons. Dark grey boxes indicate untranslated regions. Green indicates regions encoding the AAA ATPase domain. Purple indicates the region encoding the Vsp4c domain. Scale bar, 500 bp **(D)** Table of Mp*ktn* mutants including their mutated gene sequences and predicted protein sequences. K3 lines were generated with sgRNA-K3 targeting the exon 4, and K5 lines were generated with sgRNA-K5 targeting exon 7. Presented are 10 out of the 16 Mp*ktn* lines generated. **(E)** Diagram of the MpKTN protein structure and length in wild type and three selected Mp*ktn* mutants. Grey boxes indicate exons. Green regions indicate the AAA ATPase domain. Purple regions indicate the Vsp4c domain. Black regions indicate mutated sequences. **(F)** Diagram showing the generation of Cas9-free Mp*ktn* mutants expressing the microtubule reporter, *p*Mp*EF1α:GFP-*Mp*TUB1* (GFP-MpTUB1).

**Supplementary Figure 2:**
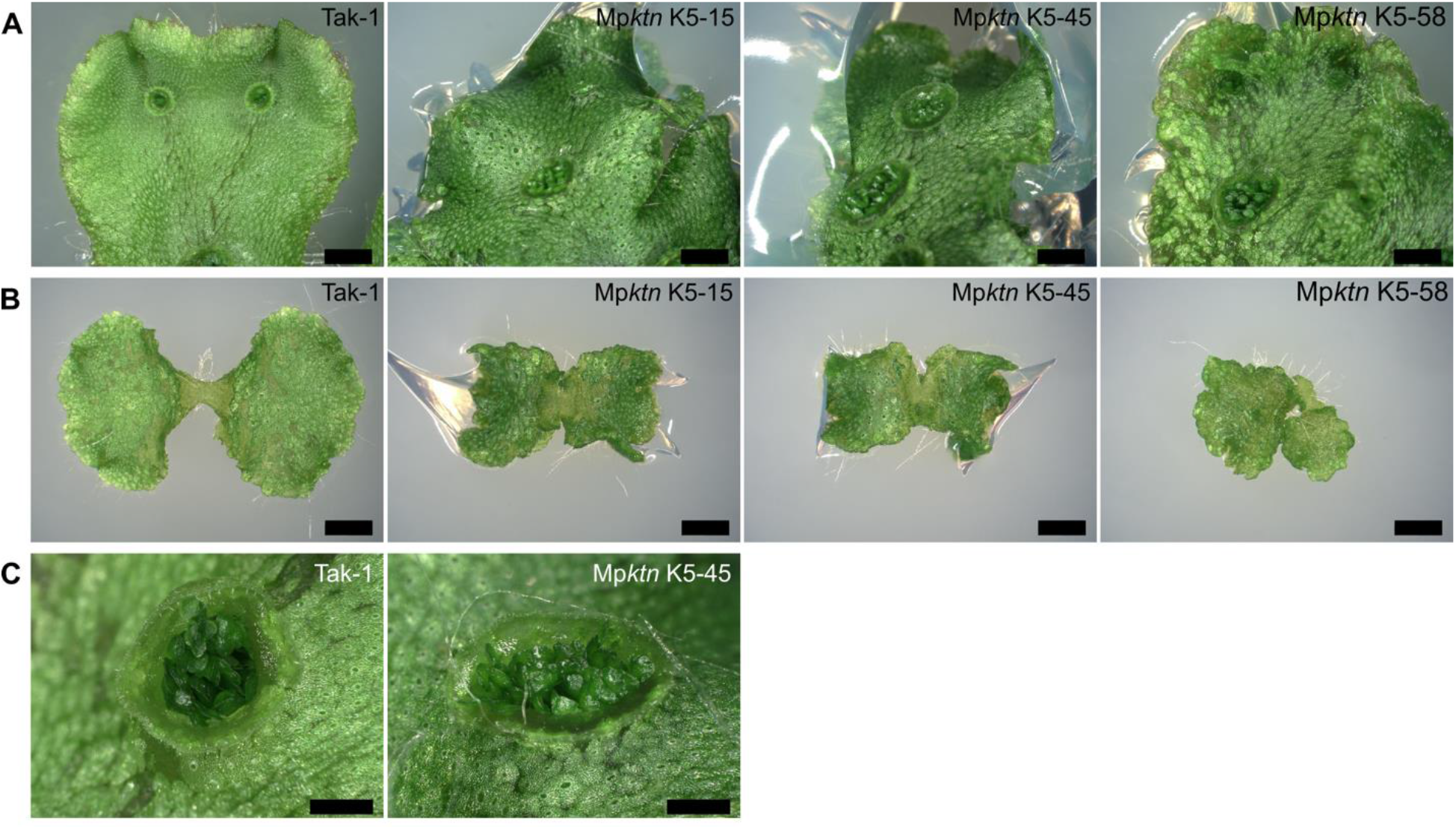
Mutations in the Mp*KTN* gene region encoding the VSP4 domain also results in defective *M. polymorpha* development. **(A - B)** Dorsal thallus (A) and 10-day-old gemmalings (B) of Tak-1 and three Mp*ktn* lines with mutations in the encoded Vsp4c domain. Scale bars, 2 mm. **(C)** Gemma cups on the dorsal thallus of Tak-1 and Mp*ktn* with a mutation in the encoded Vsp4c domain. Scale bars, 1 mm.

**Supplementary Figure 3:**
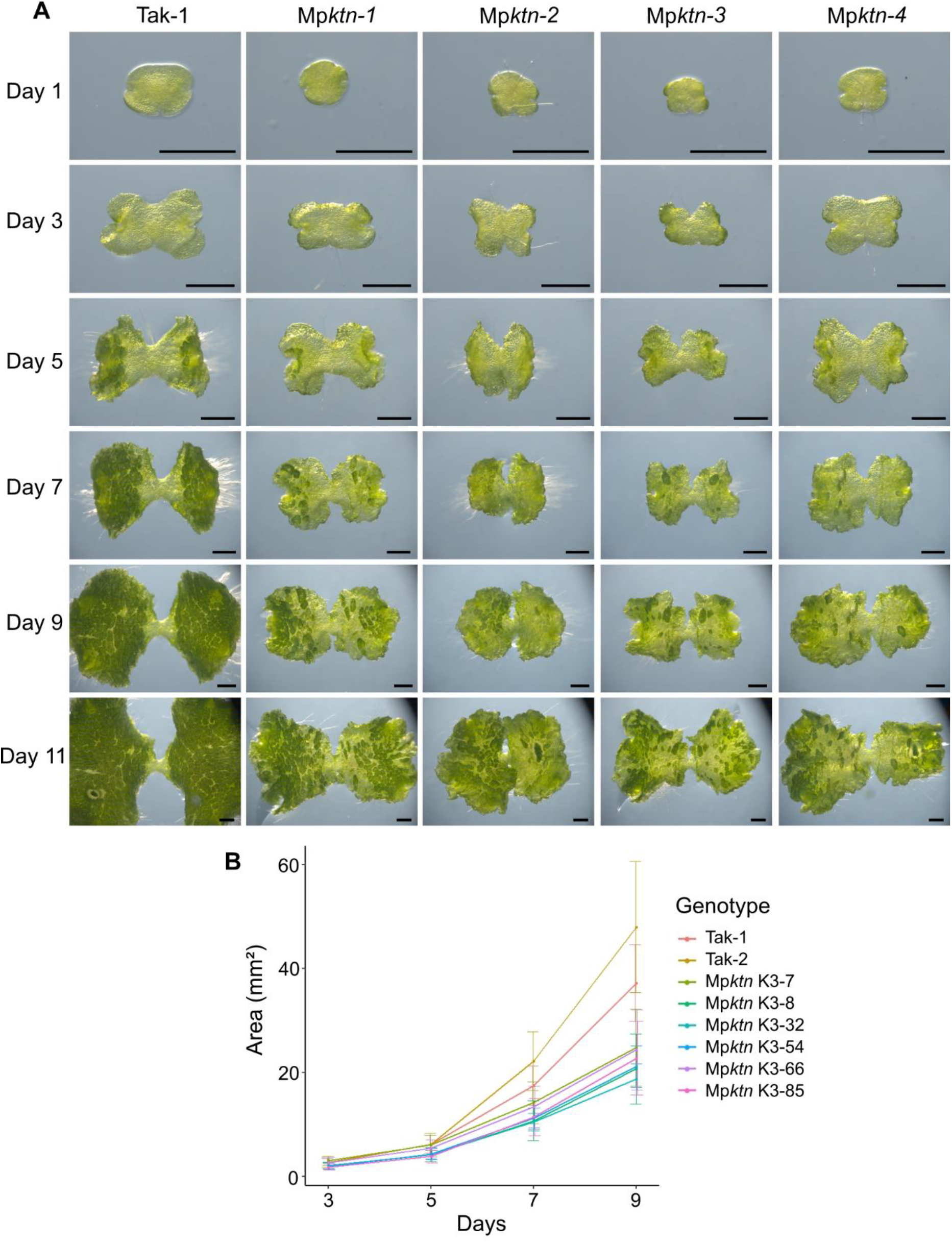
Development of Mp*ktn* gemmalings is defective and slower than wild type development. **(A)** Development of gemmalings from wild type and four Mp*ktn* lines over 11 days. Scale bars, 1 mm. **(B)** Plot of gemmaling tissue area over 9 days growth for wild type and six Mp*ktn* lines with mutations in the encoded AAA ATPase domain. Tissue area was calculated from chlorophyll autofluorescence. 9 plants per genotype were measured every 2 days. Presented is the mean area (mm²) and standard error at each timepoint.

**Supplementary Figure 4:**
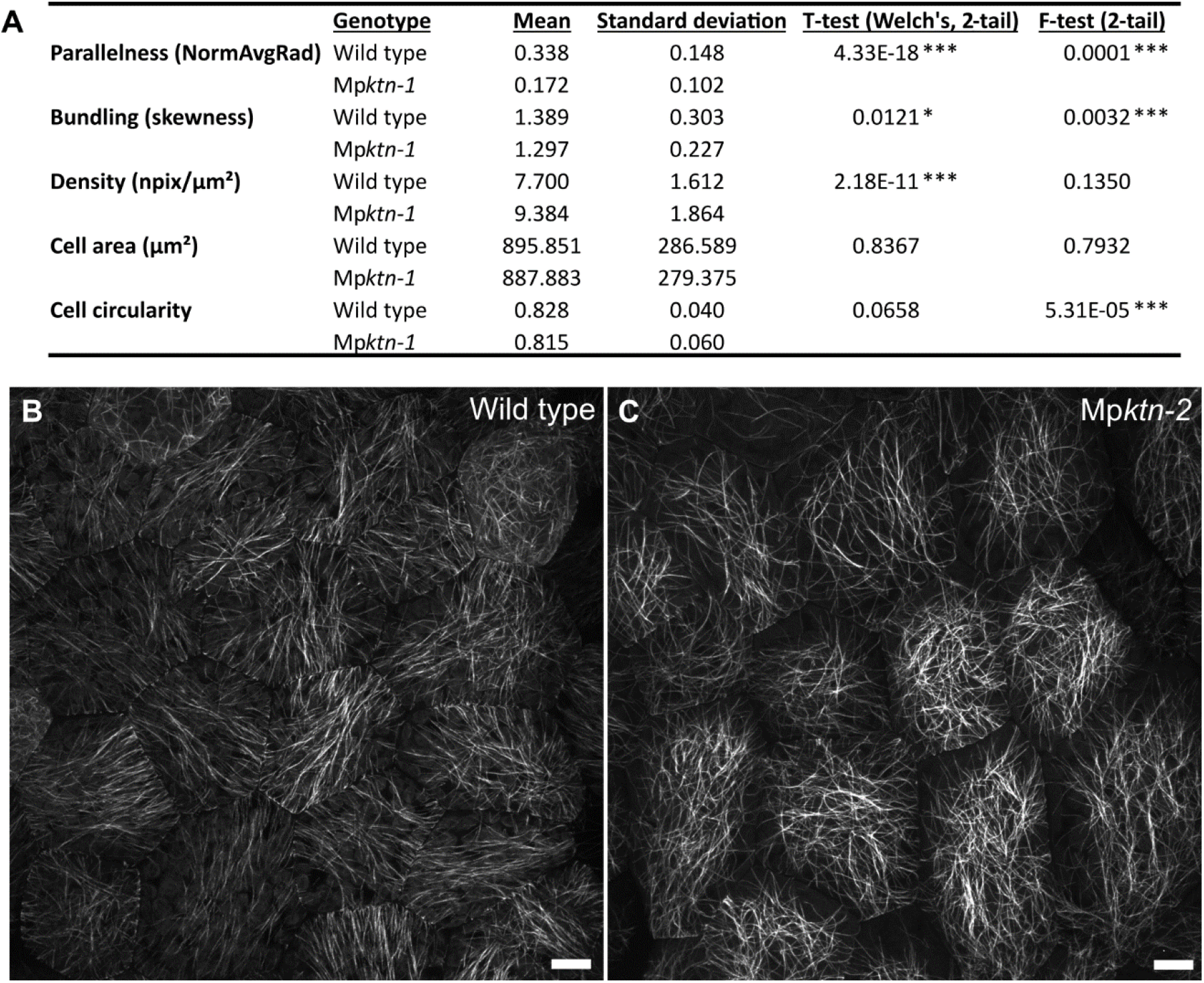
Mp*KTN* promotes parallelness and bundling of cortical microtubules. **(A)** Quantification of the parallelness, bundling and density of cortical microtubules in wild type and Mp*ktn-1* epidermal cells. N = 108 cells for wild type, N = 107 cells for Mp*ktn-1* across 10 gemmalings per genotype. Data was analysed using Welches 2-tail T-test and 2-tail F-test. Significant differences in the T-test or F-test values between the genotypes is indicated by (*) P ≤ 0.05, (**) P ≤ 0.01 and (***) P ≤ 0.001. **(B - C)** Cortical microtubule arrays in the central epidermal cells of wild type (B) and Mp*ktn-2* (C) gemmalings. Presented are Z-projections. Scale bars, 10 μm.

**Supplementary Figure 5:**
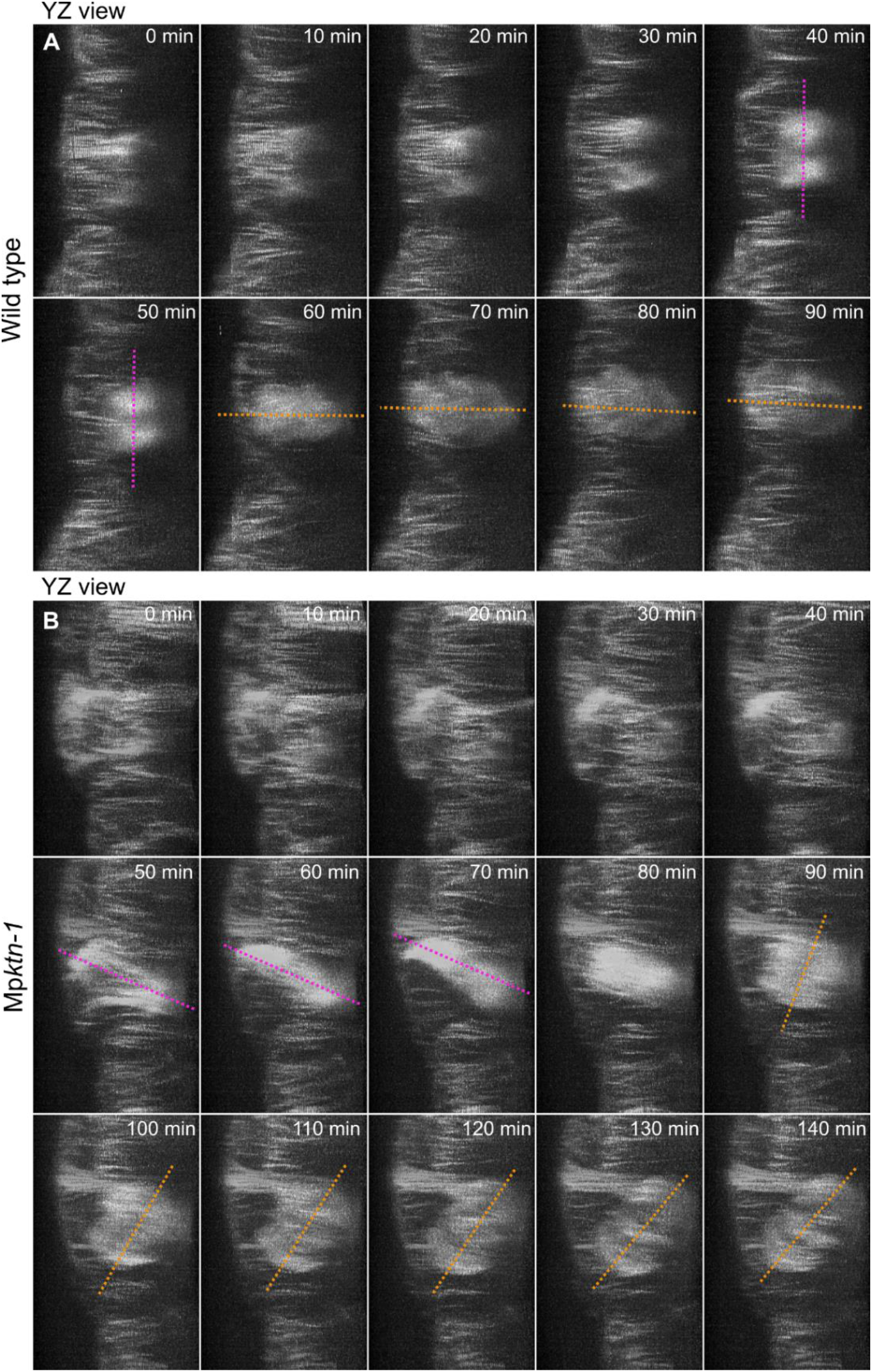
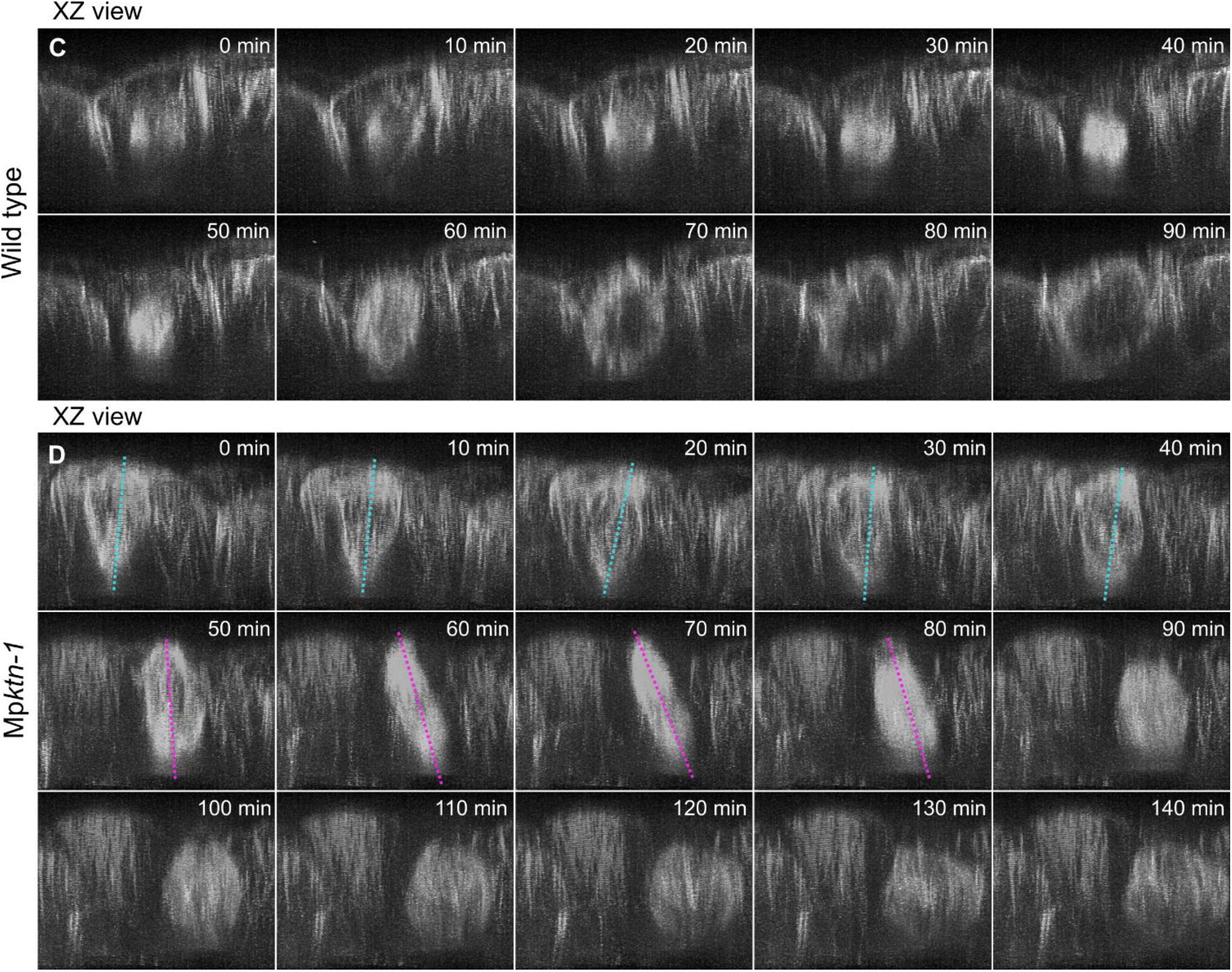
YZ and XZ views of dividing wild type and Mp*ktn-1* cells shows tilting of polar organisers, mitotic spindle and phragmoplast relative to the cell surface in Mp*ktn-1*. Timelapses of microtubule organisations in dividing wild type (**A, C**) and Mp*ktn-1* (**B, D**) cells (from Figure 4A, B) viewed from the YZ plane (A, B) and XZ plane (C, D). Presented are deconvolved Z-projections viewed from the YZ and XZ planes. Dotted cyan lines indicate the polar organiser axis, dotted magenta lines indicate the mitotic spindle axis and dotted orange lines indicate phragmoplast plane.

## References

Bettencourt-Dias, M. and Glover, D. M. (2007). Centrosome biogenesis and function: Centrosomics brings new understanding. Nat. Rev. Mol. Cell Biol. 8, 451–463.

Brown, R. C. and Lemmon, B. E. (1990). Polar organizers mark division axis prior to preprophase band formation in mitosis of the hepatic Reboulia hemisphaerica (Bryophyta). Protoplasma 156, 74–81.

Brown, R. C. and Lemmon, B. E. (2011). Dividing without centrioles: Innovative plant microtubule organizing centres organize mitotic spindles in bryophytes, the earliest extant lineages of land plants. AoB Plants 11, 1–9.

Brown, R. C., Lemmon, B. E. and Horio, T. (2004). γ-Tubulin localization changes from discrete polar organizers to anastral spindles and phragmoplasts in mitosis of Marchantia polymorpha L. Protoplasma 224, 187–193.

Buschmann, H. and Zachgo, S. (2016). The Evolution of Cell Division: From Streptophyte Algae to Land Plants. Trends Plant Sci. 21, 872–883.

Buschmann, H., Holtmannspötter, M., Borchers, A., O’Donoghue, M. T. and Zachgo, S. (2016). Microtubule dynamics of the centrosome-like polar organizers from the basal land plant Marchantia polymorpha. New Phytol. 209, 999–1013.

Chan, J., Calder, G. M., Doonan, J. H. and Lloyd, C. W. (2003). EB1 reveals mobile microtubule nucleation sites in Arabidopsis. Nat. Cell Biol. 5, 967–971.

Deinum, E. E., Tindemans, S. H., Lindeboom, J. J. and Mulder, B. M. (2017). How selective severing by katanin promotes order in the plant cortical microtubule array. Proc. Natl. Acad. Sci. U. S. A. 114, 6942–6947.

Gao, L. L., Xu, F., Jin, Z., Ying, X. Y. and Liu, J. W. (2019). Microtubule-severing protein Katanin p60 ATPase-containing subunit A-like 1 is involved in pole-based spindle organization during mouse oocyte meiosis. Mol. Med. Rep. 20, 3573–3582.

Higaki, T. (2017). Quantitative evaluation of cytoskeletal organizations by microscopic image analysis. Plant Morphol. 29, 15–21.

Higaki, T., Kutsuna, N., Sano, T., Kondo, N. and Hasezawa, S. (2010). Quantification and cluster analysis of actin cytoskeletal structures in plant cells: Role of actin bundling in stomatal movement during diurnal cycles in Arabidopsis guard cells. Plant J. 61, 156– 165.

Honkanen, S., Jones, V. A. S., Morieri, G., Champion, C., Hetherington, A. J., Kelly, S., Proust, H., Saint-Marcoux, D., Prescott, H. and Dolan, L. (2016). The Mechanism Forming the Cell Surface of Tip-Growing Rooting Cells Is Conserved among Land Plants. Curr. Biol. 26, 3238–3244.

Ishizaki, K., Chiyoda, S., Yamato, K. T. and Kohchi, T. (2008). Agrobacterium-mediated transformation of the haploid liverwort Marchantia polymorpha L., an emerging model for plant biology. Plant Cell Physiol. 49, 1084–1091.

Katoh, K., Misawa, K., Kuma, K. I. and Miyata, T. (2002). MAFFT: A novel method for rapid multiple sequence alignment based on fast Fourier transform. Nucleic Acids Res. 30, 3059–3066.

Kirchhelle, C. and Moore, I. (2017). A simple chamber for long-term confocal imaging of root and hypocotyl development. J. Vis. Exp. 1–9.

Komis, G., Luptovčiak, I., Ovečka, M., Samakovli, D., Šamajová, O. and Šamaj, J. (2017). Katanin effects on dynamics of cortical microtubules and mitotic arrays in Arabidopsis thaliana revealed by advanced live-cell imaging. Front. Plant Sci. 8, 1–19.

Kumar, S., Stecher, G., Li, M., Knyaz, C. and Tamura, K. (2018). MEGA X: Molecular evolutionary genetics analysis across computing platforms. Mol. Biol. Evol. 35, 1547– 1549.

Lei, Y., Lu, L., Liu, H. Y., Li, S., Xing, F. and Chen, L. L. (2014). CRISPR-P: A web tool for synthetic single-guide RNA design of CRISPR-system in plants. Mol. Plant 7, 1494– 1496.

Letunic, I. and Bork, P. (2018). 20 years of the SMART protein domain annotation resource. Nucleic Acids Res. 46, D493–D496.

Lindeboom, J. J., Nakamura, M., Hibbel, A., Shundyak, K., Gutierrez, R., Ketelaar, T., Emons, A. M. C., Mulder, B. M., Kirik, V. and Ehrhardt, D. W. (2013). A mechanism for reorientation of cortical microtubule arrays driven by microtubule severing. Science. 342, 1–11.

Loughlin, R., Wilbur, J. D., McNally, F. J., Nédélec, F. J. and Heald, R. (2011). Katanin contributes to interspecies spindle length scaling in Xenopus. Cell 147, 1397–1407.

Luptovčiak, I., Komis, G., Takáč, T., Ovečka, M. and Šamaj, J. (2017). Katanin: A sword cutting microtubules for cellular, developmental, and physiological purposes. Front. Plant Sci. 8, 1–10.

McNally, F. J. and Roll-Mecak, A. (2018). Microtubule-severing enzymes: From cellular functions to molecular mechanism. J. Cell Biol. 217, 4057–4069.

McNally, F. J. and Vale, R. D. (1993). Identification of katanin, an ATPase that severs and disassembles stable microtubules. Cell 75, 419–429.

McNally, K., Audhya, A., Oegema, K. and McNally, F. J. (2006). Katanin controls mitotic and meiotic spindle length. J. Cell Biol. 175, 881–891.

Rasmussen, C. G., Wright, A. J. and Müller, S. (2013). The role of the cytoskeleton and associated proteins in determination of the plant cell division plane. Plant J. 75, 258– 269.

Sauret-Güeto, S., Frangedakis, E., Silvestri, L., Rebmann, M., Tomaselli, M., Markel, K., Delmans, M., West, A., Patron, N. J. and Haseloff, J. (2020). Systematic Tools for Reprogramming Plant Gene Expression in a Simple Model, Marchantia polymorpha. ACS Synth. Biol. 9, 864–882.

Schindelin, J., Arganda-Carreras, I., Frise, E., Kaynig, V., Longair, M., Pietzsch, T., Preibisch, S., Rueden, C., Saalfeld, S., Schmid, B., et al. (2012). Fiji: An open-source platform for biological-image analysis. Nat. Methods 9, 676–682.

Sugano, S. S., Shirakawa, M., Takagi, J., Matsuda, Y., Shimada, T., Hara-Nishimura, I. and Kohchi, T. (2014). CRISPR/Cas9-mediated targeted mutagenesis in the liverwort Marchantia polymorpha L. Plant Cell Physiol. 55, 475–481.

Thamm, A., Saunders, T. E. and Dolan, L. (2020). MpFEW RHIZOIDS1 miRNA-Mediated Lateral Inhibition Controls Rhizoid Cell Patterning in Marchantia polymorpha. Curr. Biol. 30, 1905–1915.

Zhang, Q., Fishel, E., Bertroche, T. and Dixit, R. (2013). Microtubule severing at crossover sites by katanin generates ordered cortical microtubule arrays in Arabidopsis. Curr. Biol. 23, 2191–2195.

